# Rethinking causality and data complexity in brain lesion-behaviour inference and its implications for lesion-behaviour modelling

**DOI:** 10.1101/2019.12.17.878355

**Authors:** Christoph Sperber

## Abstract

Modelling behavioural deficits based on structural lesion imaging is a popular approach to map functions in the human brain, and efforts to translationally apply lesion-behaviour modelling to predict post-stroke outcomes are on the rise. The high-dimensional complexity of lesion data, however, evokes challenges in both lesion behaviour mapping and post stroke outcome prediction. This paper aims to deepen the understanding of this complexity by reframing it from the perspective of causal and non-causal dependencies in the data, and by discussing what this complexity implies for different data modelling approaches. By means of theoretical discussion and empirical examination, several common strategies and views are challenged, and future research perspectives are outlined. A main conclusion is that lesion-behaviour inference is subject to a lesion-anatomical bias that cannot be overcome by using multivariate models or any other algorithm that is blind to causality behind relations in the data. This affects the validity of lesion behaviour mapping and might even wrongfully identify paradoxical effects of lesion-induced functional facilitation – but, as this paper argues, only to a minor degree. Thus, multivariate lesion-brain inference appears to be a valuable tool to deepen our understanding of the human brain, but only because it takes into account the functional relation between brain areas. The perspective of causality and inter-variable dependence is further used to point out challenges in improving lesion behaviour models. Firstly, the dependencies in the data open up different possible strategies of data reduction, and considering those might improve post-stroke outcome prediction. Secondly, the role of non-topographical causal predictors of post stroke behaviour is discussed. The present article argues that, given these predictors, different strategies are required in the evaluation of model quality in lesion behaviour mapping and post stroke outcome prediction.

## 1 Introduction

One of the main objectives of cognitive neuroscience is to understand the relationship between brain anatomy and behavioural functions. A powerful method to achieve this is to study the relation between focal brain damage and behavioural deficits. Not only is lesion-behaviour mapping in cerebral stroke the oldest method in this field, it is a driving force in human brain mapping (Rorden & Karnath, 2004). In voxel-based lesion behaviour mapping (VLBM; Bates et al., 2003; Rorden, Karnath, & Bonilha, 2007), the relation between structural topographical lesion data and a behavioural target variable is statistically modelled to map functions onto the brain. Such lesion behaviour models are also clinically applicable if they are able to accurately predict performance when modelling chronic post-stroke outcomes based on acute lesion imaging.

Lesion behaviour modelling, however, is not a trivial task. A researcher can choose between a plethora of univariate and multivariate algorithms. Many strategies on how to select or dimensionally reduce features from brain imaging exist. And last but not least, evaluating or interpreting lesion-behaviour models can be challenging, especially as different applications of lesion behaviour models have different requirements and can be influenced by the choice of different parameter settings. All these challenges are magnified by the fact that lesion data are high-dimensional and complex. Lesion imaging provides us with information in a large amount of imaging voxels, i.e. the smallest units of a brain image, and the information between these voxels is not independent.

The present paper aims to provide the reader with a new perspective on lesion-behaviour data and dependencies between variables in this context. With this perspective, we are able to re-evaluate present approaches in lesion behaviour modelling, and we can identify strategies to improve it. This is achieved by theoretical considerations and supported by empirical examples based on simulations using real lesion data. The present paper claims that some common methodological beliefs in the scientific field are wrong and that some commonly used analysis strategies are limited in their current implementation.

## 2 General Methods and data availability

All empirical examples in this paper are based on a publicly available dataset of 131 normalised left hemisphere stroke lesions from the Moss Rehabilitation Research Institute at the University of Pennsylvania, as available in the LESYMAP software (https://github.com/dorianps/LESYMAP). This dataset contains binary lesion maps obtained from either CT or structural MRI imaging that were normalised to MNI-space (see Pustina, Avants, Faseyitan, Medaglia, & Coslett, 2018 for further information). Analyses were performed with Matlab 2018a, SPM12 (https://www.fil.ion.ucl.ac.uk/spm/), libSVM 3.23 (Chang & Lin, 2011), and NiiStat (https://www.nitrc.org/projects/niistat/). Topographies were visualised using MRIcron (https://www.nitrc.org/projects/mricron) and MRIcroGL (https://www.nitrc.org/projects/mricrogl). Several empirical examples in this paper are based on simulated behavioural variables (for details and rationale see Mah, Husain, Rees, & Nachev, 2014; Pustina et al., 2018; Sperber & Karnath, 2018). Such simulations are a powerful tool to validate many aspects of lesion behaviour mapping methods. However, there is no set of perfectly realistic simulation parameters, and thus ecological validity is limited. All simulated data underlying shown examples, simulation scripts, lesion overlay maps, and statistical topographies are publicly available at https://data.mendeley.com/datasets/ydm2rfk9sn/2. No part of the study procedures or analyses was pre-registered in a time-stamped, institutional registry prior to the research being conducted.

## 3 Causality and correlation in lesion-behaviour relationships

Lesion-behaviour mapping is commonly seen as a method that – contrary to methods such as fMRI, EEG, or MEG – allows us to draw conclusions about *causal* relationships between behaviour and brain (Rorden & Karnath, 2004; Zavaglia et al., 2015). That means that if we identify brain regions that are associated with a deficit in lesion-behaviour mapping, we can assume that the region’s functionality is essential, or, in other words, *causal* for the behaviour. Voxel-based lesion behaviour mapping (Bates et al., 2003; Rorden et al., 2007), a mass-univariate brain mapping method based on the framework of statistical parametric mapping (Friston et al., 1995), has been widely used and is the most popular method following this rationale of brain lesion-behaviour inference.

After several years of flourishing success, the popularity and authority of VLBM was suddenly overshadowed by findings of systematic errors in the VLBM method (Mah et al., 2014). Using simulation approaches, Mah and colleagues showed that topographical results were misplaced towards the centres of the major cerebral arteries. Although some methodological objections against the size of the bias and its operationalisation exist (Sperber & Karnath, 2017; Pustina et al., 2018; Sperber, Wiesen, & Karnath, 2019a), several studies replicated this finding (Inoue, Madhyastha, Rudrauf, Mehta, & Grabowski, 2014; Sperber & Karnath, 2017; Sperber et al., 2019a), suggesting that a bias of statistical topographies is indeed an issue inherent to the VLBM method.

This limitation of VLBM was attributed to its mass-univariate design, i.e. the fact that in VLBM each voxel is tested independently of all other voxels in the brain. This contrasts with actual brain anatomy in two ways: voxels in the brain are 1) lesion-anatomically dependent and 2) functionally dependent. Firstly, the voxel-voxel lesion status, i.e. how regularly voxels are damaged together in individual patients, is systematic due to lesion anatomy, i.e. due to stroke following the vasculature and arteries’ varying susceptibility to stroke. This violation of independence is thought to induce the topographical bias along the vasculature (Mah et al., 2014; Sperber & Karnath, 2017; Karnath, Sperber, & Rorden, 2018). Secondly, brain functions are not anatomically organized in single voxels, but in larger areas or networks. The violation of functional dependence induces the so-called partial injury problem (Rorden, Fridriksson, & Karnath, 2009; Sperber et al., 2019a), which leads to reduced statistical power when larger functional modules are often only damaged in parts. Accordingly, it has been shown that VLBM often fails in simulation settings where two or more distinct brain areas underpin a function (Mah et al., 2014; Zhang, Kimberg, Coslett, Schwartz, & Wang, 2014; Pustina et al., 2018) by missing some, or all, areas.

The impact of these findings was intensified by a parallel methodological innovation – the establishment of multivariate lesion behaviour mapping (MLBM) methods. These methods allow for mapping lesion behaviour inference by using computational models that account for the status of all voxels at once (e.g. Mah et al., 2014; Zhang et al., 2014; Pustina et al., 2018). It has been postulated that MLBM, which accounts for the mutual dependence of voxels, is the solution to the limitations of mass-univariate VLBM, including the lesion-anatomical dependence (Mah et al., 2014; Xu, Jha, & Nachev, 2018). Large parts of the lesion behaviour mapping community adapted this view and repeated the claim that MLBM resolves these limitations of VLBM (e.g., Adolphs, 2016; Carter et al., 2017; Toba et al., 2017; DeMarco & Turkeltaub, 2018; Thye et al., 2018; Xu et al., 2018; Zhao et al., 2018; Dickens et al., 2019; Gläscher, Adolphs, & Tranel, 2019; Howard, Smith, Coslett, Buxbaum, & Krakauer, 2019; Valero-Cabré, Toba, Hilgetag, & Rushmore, 2019; Wong, Jax, Smith, & Buxbaum, 2019). However, empirical findings do not fully support this view. Indeed, it has been shown that MLBM is often superior to VLBM in identifying brain modules that consist of multiple regions or networks (Zhang et al., 2014; Pustina et al., 2018; Sperber, Wiesen, Goldenberg, & Karnath, 2019b). Thus, MLBM seems to resolve the partial injury problem. On the other hand, so far it has not been shown that the topographical bias is absent in MLBM. Instead, a lesion-anatomically induced bias of topographical results comparable to the bias in VLBM was found for an MLBM method based on support vector regression (Sperber et al., 2019a). Still, it has been claimed that the main reason to use multivariate instead of mass-univariate methods is that MLBM accounts for lesion-anatomical dependence and resolves the issue of topographical bias, and therefore univariate methods must be replaced by multivariate methods (Nachev, 2015; Xu et al., 2018).

To sum up, several studies claim that MLBM solves the limitations of mass-univariate lesion behaviour, and large parts of the field embraced this view. This position stands vis-à-vis the empirical finding that at least one MLBM method based on support vector regression does not live up to this expectation (Sperber et al., 2019a). One can argue that the reason might be shortcomings of just this MLBM method, but not an issue of MLBM in general. This paper argues that the inability to account for lesion-anatomical dependence is indeed a shortcoming of *any* common multivariate approach to lesion behaviour mapping. To strengthen this point, it does not suffice to empirically evaluate some MLBM methods. Rather, it requires strong, universal theoretical arguments. These can be found by reframing the term ‘causality’ in lesion behaviour mapping.

### 3.1 Rethinking lesion behaviour modelling with causal models

Before the scientific field was aware of topographical biases in VLBM, a common view was that significant findings in VLBM generally indicated regions in which damage is causal for a deficit. The concept of causality can intuitively be illustrated with graphical causal models (see Pearl, Glymour, & Jewell, 2016). If two adjacent significant regions A and B were found, the classical VLBM perspective assumed damage to each feature to be causal for the deficit individually (Fig.1A). This view was challenged by the discovery of the topographical bias, meaning that in our example damage to region B might indeed not be causal for the deficit, although statistically significant voxels are found there (Fig.1B). This can be explained by taking into account the vasculature in our causal model (Fig.1C). If we assume that both regions are supplied by the same minor cerebral artery, then a stroke in this artery (or any major artery supplying this minor artery) will likely *cause* damage to both areas. The upper three nodes of the graph (stroke to the minor cerebral artery and damage to areas A and B) now constitute a fork. For such fork in graphical causal models where X is a common cause of variables Y and Z, variables Y and Z are likely dependent (see chapter 2.2. and SCM 2.2.6. in Pearl et al., 2016), or, using terminology from inferential statistics, both variables are *associated*. Thus in our example, the damage status of areas A and B will be dependent, which is the lesion-anatomical dependence between brain regions/voxels introduced in the previous section. Now, the most important point is that i) the behavioural deficit and damage to area A are causally related and dependent and that ii) damage to area A and area B are dependent. From this it follows that iii) also damage to area B and the deficit will be dependent, but without a causal relation between both. The possible dependence between damage to area B and the deficit is a mere association. Statistical tests, however, are ignorant of the concept of causality. They merely identify associations and, therefore, might also associate region B with the behavioural deficit. This is why VLBM can suffer from a topographical bias. But we can even go beyond regions A and B and look at all areas in the brain. Many areas in the brain might share the same major or minor cerebral artery with area A, which can be illustrated by an oversimplified graphical causal model (Fig. 1D). Therefore, the damage status of many regions will be dependent on the damage status of area A, and the deficit will, to some degree, be associated with many or, even all, areas in the brain. This results in statistical parametric maps that contain more or less high non-zero values in all areas of the brain, even when only damage to a small circumscribed region is causal for a behavioural deficit. To conclude, we can reframe the term ‘causality’ in lesion-behaviour modelling: brain-behaviour associations are commonly found all over the brain, and only some of them in fact reveal a causal relation. Although disease-informed brain mapping in theory allows us to draw conclusions about *causal* brain-behaviour relations, VLBM does not necessarily identify the areas where damage is causal for a deficit, but first and foremost areas with the strongest lesion-deficit associations. These will likely correspond to a high degree to the regions that are actually causal, but – as we have seen in the previous section on limitations of VLBM – with some systematic deviations.

**Figure 1.**
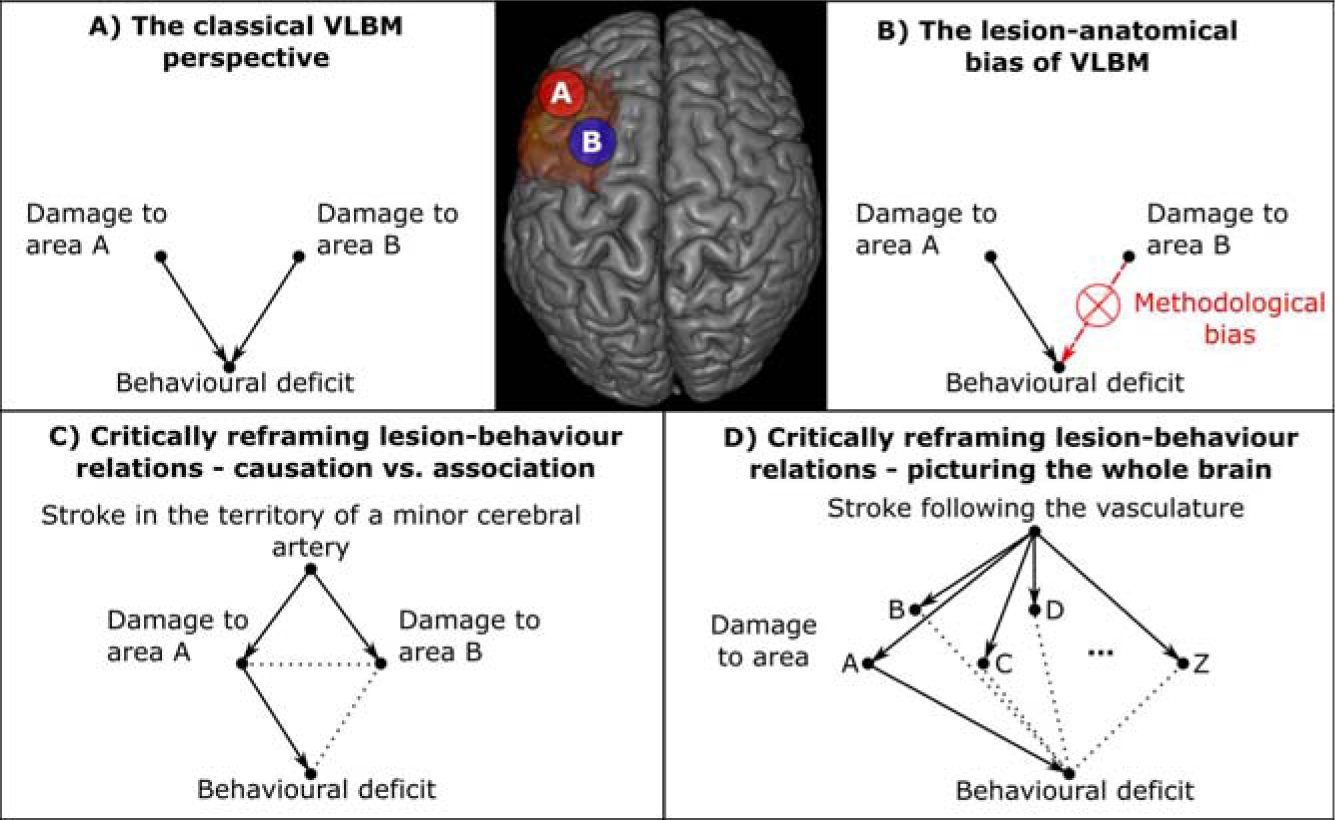
Graphical causal models of lesion-behaviour relationships. Graphical causal models illustrate different views on causality in lesion-behaviour relationships. Nodes illustrate different variables, and arrows indicate (putative) causal relations between these variables. Dotted lines indicate that no causal relation between variables, but merely a non-causal association exists. The brain image is a statistical map taken from the VLBM analysis seen in Figure 3, based on a simulation with the pars triangularis as ground truth. To keep the illustration simple, the non-causal associations between damage in different areas in panel D are not shown.

With this new perspective of causality in lesion behaviour relations, what do we learn about the (in)ability of multivariate lesion behaviour mapping to overcome the systematic deviations of VLBM? How can any lesion-brain inference method resolve the issues raised by lesion-anatomical dependence between voxels? The answer is: the method needs to draw *causal* inference. If, and only if, a multivariate inference method can separate between a mere (non-causal) association and a causal relation, it will not suffer from limitations originating from lesion-anatomical dependence.

Are multivariate approaches, such as multiple regression or machine learning algorithms using support vector machines, support vector regression, or neural networks, able to draw causal inference? No, they are not. “Machine learning programs (including those with deep neural networks) operate almost entirely in an associational mode. They […] attempt to fit a function, in much the same way that a statistician tries to fit a line to a collection of points.” (see p.30 in Pearl & Mackenzie, 2018). The only way to draw causal inference with machine learning algorithms or classical inferential statistics is a randomised controlled experimental design, where the researcher manipulates the independent variable. This is not the case in lesion-behaviour data, and thus multivariate methods are not the solution to all limitations of VLBM.

### 3.2 Simulation example

If MLBM supposedly still suffers from limitations due to lesion-anatomical dependence, it remains an open question why MLBM using a support-vector machine (SVM) performed so well in the second experiment of the study by Mah et al. (2014). In this study they investigated a simulation based on damage to the gyrus angularis (Brodmann area 39) and parts of the inferior frontal gyrus (Brodmann areas 44). In this example, mirroring different conflicting theories about the neural correlates of spatial neglect, VLBM grossly misplaced parts of the topographical results into the superior temporal gyrus. On the other hand, the voxels that contributed the most to the multivariate lesion-behaviour model using a linear SVM were situated in the actual ground truth areas. The error encountered in VLBM in the latter example does not resemble the above-mentioned description of the partial injury problem, where some areas might simply not be correctly identified by VLBM due to decreased statistical power. Significant voxels were found in an entirely different area, suggesting that a spatial lesion-anatomical bias was responsible for the incorrect results. If this is truly the case, then MLBM by SVM might have, in fact, overcome the issues originating from lesion-anatomical dependence. However, as we will see, these findings were only a part of the picture.

To critically re-evaluate this previous example from the study by Mah and colleagues, I replicated the simulation experiment on the example lesion data of 131 left hemisphere stroke patients. The same parameters were chosen, i.e. the behavioural deficit was simulated based on damage to the ‘ground truth’ areas 39 and 44 of the Brodmann atlas in MRIcron (Fig. 2A). If more than 20% of voxels in either area were damaged, a patient was considered to suffer from a deficit with a probability of 90%. The deficit was thus a binary one and lesion-behaviour relations were analysed in each voxel affected in at least 5 patients using mass-univariate Liebermeister-tests with Bonferroni-correction at p < 0.05 as implemented in NiiStat. Note that the Liebermeister test is a common binomial test in VLBM, and it only differs from Fisher’s exact test in minor details (Rorden et al., 2007).

**Figure 2.**
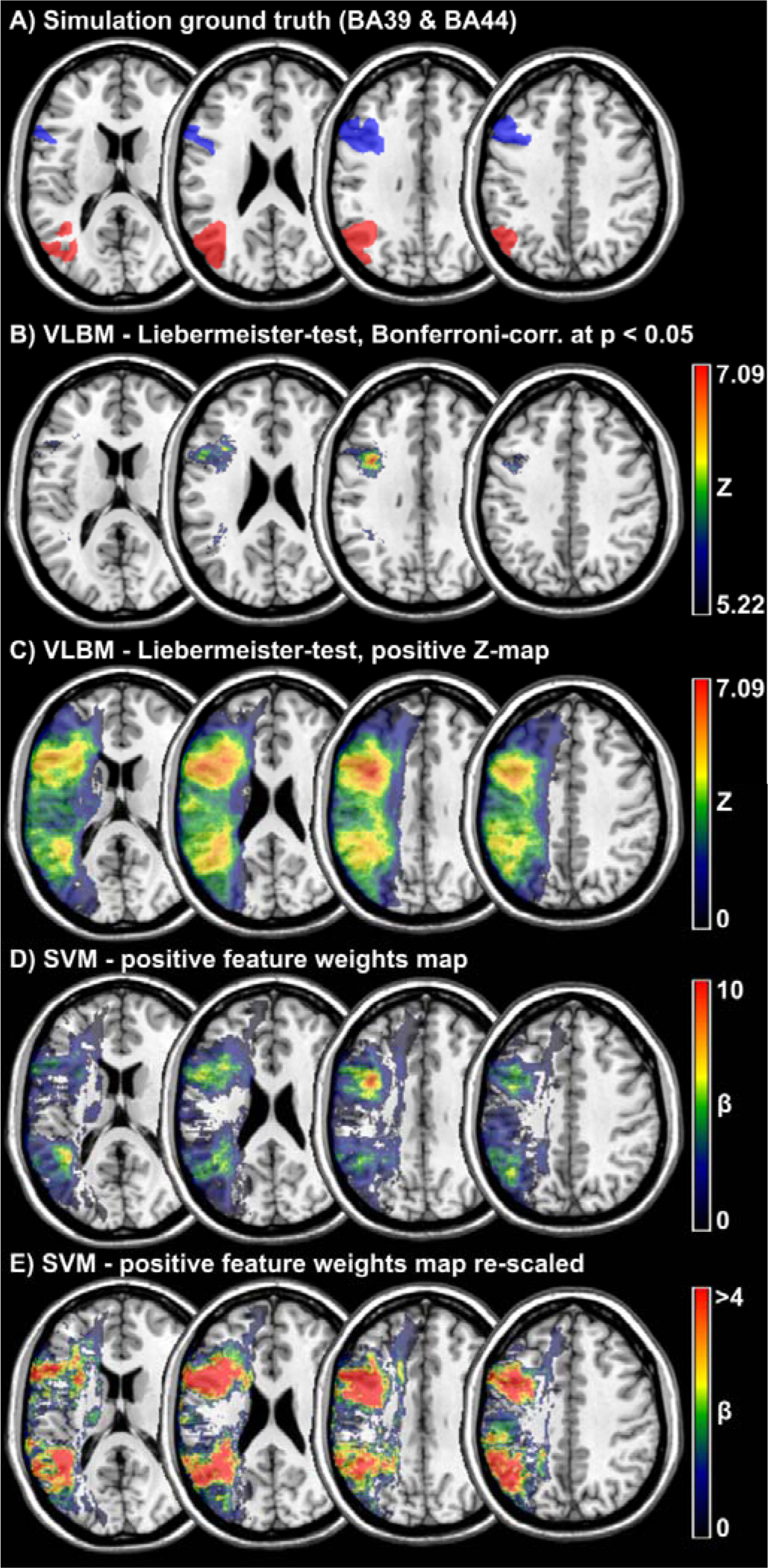
Comparison of mass-univariate and multivariate lesion-brain inference. A) In the example lesion data set, a behavioural symptom was simulated for each patient based on damage to Brodmann areas 39 and 44. B) and C) Lesion behaviour mapping of these simulated behavioural data was first performed with a mass-univariate VLBM by the Liebermeister test (Bonferroni correction threshold for p < .05 at Z = 5.22, maximum Z = 7.09). D) and E) The same data were also mapped using a high-dimensional approach based on a support vector machine. Feature weights were scaled to a maximum value of 10. Note that negative feature weights are not shown.

The results were highly unexpected in one way: no misplacement of statistical results into the superior temporal gyrus was found (Fig. 2B). Most of the 8719 significant voxels were located in or close to Brodmann area 44; whereas, only a small number of significant voxels were found in or around Brodmann area 39. A typical example of the partial injury problem can be seen in the present simulation, with reduced statistical power to identify all areas in a multi-region module. Visual inspection of the topography reveals a slight shift of significant clusters away from the cortex towards the centre of the territorial artery. However, no gross misplacement of the results was found. However, investigation of the non-thresholded Z-map (Fig. 2C) revealed that there were indeed several sub-threshold peaks in brain areas located between both Brodmann areas, including subcortical and temporal areas (Fig. 2C). Thus, although the results did not exactly replicate the findings by Mah and colleagues, the present results still find enlarged signal in temporal areas to some degree. The discrepancies between both studies likely arose from differences in sample characteristics, which have been shown to affect results in VLBM (Gajardo-Vidal et al., 2018; Sperber & Karnath, 2018).

Let us now look at this example and how functional and lesion-anatomical dependence between voxels induced errors in the VLBM analysis. Firstly, both Brodmann areas are functionally dependent. Therefore, the partial injury problem reduces statistical power in the areas where we want the VLBM analysis to produce significant results. This was the case in Brodmann area 39 in the present example and, to some degree, also in both areas in the former experiment by Mah and colleagues. Note that in their online available supplementary 3D figure it becomes apparent that a majority of significant voxels in their topography were still also found in, or neighbouring, the ground truth areas, and only a smaller proportion was misplaced into the superior temporal gyrus. Secondly, the lesion-anatomical dependence lead to an association of subcortical, as well as, temporal areas with the symptom. This happens in brain areas where damage often coincides with damage to ground truth areas. In this example, the association is especially high as temporal areas might often be damaged together with either Brodmann area 39 or 44.

Is high-dimensional, multivariate lesion brain inference able to overcome these issues? To answer this question, the same data were also analysed with MLBM based on a linear kernel SVM, mirroring the analysis by Mah and colleagues. To do so, the SVR-LSM Toolbox (Zhang et al., 2014) was modified and a hyper-parameter optimisation was added. All voxels damaged in at least 5 patients were included as features to predict the simulated behavioural score. Hyper-parameter C was optimised by a 10-fold cross-validation for C ranging from 2^−20^ to 2^20^. Model fit was maximal for C = 2^−16^ with a cross-validation prediction accuracy of 89.3%. Individual feature weights were then scaled to a maximum of 10 and re-mapped into brain space. The SVM feature-weight-map (Fig. 2D) demonstrated peak values in both Brodmann areas, which resembled the significant areas of the VLBM (Fig. 2B). A visual comparison with the full VLBM Z-map showed that peaks in the SVM feature weights more clearly stood out of the signal in the surrounding areas. A closer look at the lower feature weight range (Fig. 2E), however, revealed several aspects: i) areas with larger signal in the VLBM Z-map (Fig. 2C) and SVM feature weight map (Fig. 2E) appeared to be relatively similar, but with the VLBM Z-map having much smoother transitions between high and low signal areas, ii) areas with larger feature-weights appeared to be slightly shifted away from cortical ground truth areas towards the centre of the arterial territory in a similar manner as areas with larger Z-values, and iii) a few smaller clusters with larger feature-weights were scattered across areas situated between both Brodmann areas, for example, in subcortical and temporal areas.

In summary, visual comparison of the VLBM Z-map and the SVM feature weights-map indicates that, in this example, the multivariate model indeed outperforms mass-univariate models as it provides a stronger and more clear-cut signal in the ground truth areas. Still, the SVM did not only utilise information in voxels that were *causally* linked to the deficit through the simulation, but also in voxels that were non-causally associated with the symptom in voxels neighbouring the ground truth areas, and occasionally even in areas further away. Doing so, the map of these voxels bore some resemblance to the VLBM map.

Thus, it seems that multivariate inference is not hampered by functional dependence between areas, but still by lesion-anatomical dependence. Not affected by functional dependence and the partial injury problem, the SVM in the previous study by Mah and colleagues – contrary to VLBM – correctly found peak values in the ground truth regions. At the same time, the SVM likely also used information from temporal areas. The reason for this is that damage in these temporal areas was associated with the symptom and helpful in the prediction. However, due to the multivariate approach accounting for functional dependence, information from temporal areas was less helpful in the prediction than information from the ground truth areas in frontal and parietal cortex, and therefore peak voxels were found in the ground truth areas.

### 3.3. Conclusion

High-dimensional, multivariate lesion-brain inference is a valuable innovation that allows us to advance our knowledge about functional brain anatomy, and it has the potential to replace former mass-univariate methods in parts or even entirely. Nevertheless, it is not perfect. The high-dimensional complexity of anatomical lesion data results in a situation where only causal inference can yield perfectly non-biased results, and such causal inference is not employable in a hypothesis-free, high-dimensional inference problem (see Pearl et al., 2016). Thus, it seems that we have to accept that lesion-behaviour mapping, just like any other method in neuroscience, is imperfect.

A last word on post stroke outcome prediction. The lesion-anatomical bias has been shortly discussed in this context (Price, Hope, & Seghier, 2017), but is it indeed relevant? Commonly, multivariate machine learning algorithms are used in this field (e.g. Rondina, Filippone, Girolami, & Ward, 2016; Rondina, Park, & Ward, 2017; Hope, Leff, & Price, 2018; Loughnan et al., in press), and, therefore, no difference is made between features that are causally linked to or only non-causally associated with the target variable. Both might be used to increase the predictive performance of these models, and if we only follow a pragmatic approach, this should not bother us. If we can obtain a good prediction of lung cancer occurrence by looking at the number of work breaks somebody takes or from knowing if a person carries a lighter in their pockets, this can be clinically useful, even if carrying household objects in your pocket is no cause for cancer. A relevant question, however, is how algorithms deal with redundant information (e.g. smoking and carrying a lighter), which will be a topic in section 5 on feature reduction. The perspective on causality and association in lesion-behaviour data, however, comes with one important implication for stroke outcome prediction: we might find useful features for prediction outside of brain areas that are causally linked with the behaviour. For example, primary motor deficits might be predictable from topographical information taken from outside the motor system. Further, the predictive value of individual features also depends on lesion anatomy. If typical lesion anatomy differs between samples – e.g. due to different inclusion criteria with regards to maximum symptom severity, aetiology, or lesion size – the association strength between individual features and the target variable might differ. Thus, model generalisability might be limited between different clinical samples when recruitment strategies differ.

## 4 Inverse lesion-behaviour relations – paradoxical functional facilitation?

A typical consequence of cerebral damage are deficits across a wide range of cognitive, perceptual, and behavioural domains. A by far less common finding is the opposite effect – *improved* functions due to focal brain damage. A popular example is the ‘Sprague Effect’ in cats (Sprague, 1966): after unilateral lesions to all cortical visual areas, cats suffer from homonymous hemianopia and are unable to visually orient to stimuli in the contralateral hemifield. However, when the cats’ contralateral superior colliculus is also subsequently lesioned, a surprising restoration of orienting behaviour towards the contralateral hemifield can be observed. While this and most other paradoxical behavioural facilitation effects were described after experimental lesions in animals, some are also discussed in humans (for review see Kapur, 1996). For example, possibly analogous effects to the Sprague-effect may exist in humans (Fecteau, Pascual-Leone, & Théoret, 2006; Valero-Cabré et al., 2019), and enhancing facilitating effects were reported for aphasic patients in lie detection (Etcoff, Ekman, Magee, & Frank, 2000) and patients with frontal lobe damage in an arithmetic problem solving task (Reverberi, Toraldo, D’Agostini, & Skrap, 2005).

Although these cases are rare, they display exciting findings that can expand our understanding of how the brain functions (Kapur, 1996). Hence, some lesion behaviour mapping tools investigate both positive and negative lesion-behaviour relations (e.g. SVR-LSM toolbox by Zhang et al., 2014; LESYMAP by Pustina et al., 2018), and multivariate lesion behaviour mapping methods have been postulated to be of use in the investigation of such effects (Toba et al., 2017; Valero-Cabré et al., 2019). While this is surely the case, caution might be advised. If there are non-causal *positive* lesion-behaviour relations when regions are systematically damaged together, there might also be non-causal *negative* lesion-behaviour relations when regions are systematically not damaged together. To illustrate paradoxical effects by a real-world example, imagine the entirety of all current English Premier League football players. In this group, training goalkeeping will be highly predictive of low (i.e. below average) dribbling and finishing skills, i.e. goalkeeper training will negatively correlate with these field players’ skills. Does this mean that goalkeeper training causes an impairment of these skills? No. The obvious reason is that the player role of being a goalkeeper/field player is highly systematic and mutually exclusive, i.e. not being a goalkeeper who trains goalkeepers’ skills means being a field player who trains field players’ skills. For this reason, a non-causal paradoxical association exists in this example.

### 4.1 Simulation example

The emergence of non-causal negative lesion-behaviour relations in lesion-behaviour modelling can be investigated in a simulation setting in which we a priori decide to not include any causal negative relation. In the example lesion sample, a behavioural deficit was simulated based on the damage to the pars triangularis (as defined by the AAL atlas, Tzourio-Mazoyer et al., 2002). Thus, the only causal lesion-behaviour relation was a positive association between the deficit and damage to the pars triangularis. Following a previous study, the deficit was simulated based on 70% true signal and 30% uniform noise (Pustina et al., 2018) in order to obtain realistic effect sizes. Lesion behaviour mapping was performed in all voxels damaged in at least 5 patients by multivariate analysis with the SVR-LSM toolbox (Zhang et al., 2014) using direct total lesion volume control and 1000 permutations, and by univariate analysis with NiiStat with, or without, control for lesion size by nuisance regression.

As shown in Figure 3, all analyses unsurprisingly found the strongest lesion-behaviour associations in frontal regions, where lesion and behaviour were associated positively. Nonetheless, all analyses also found regions where behaviour was negatively associated with the deficit, i.e. regions in which damage was associated with less severe deficits. Negative associations were mainly found in parietal, parieto-occipital, and posterior temporal regions – regions situated far away from the inferior frontal lobe, and thus only rarely affected by stroke together with the pars triangularis. This negative signal even surpassed an uncorrected p < 0.05 in all analyses. However, it did not survive a correction for multiple comparisons by false discovery rate correction at q = 0.05.

**Figure 3.**
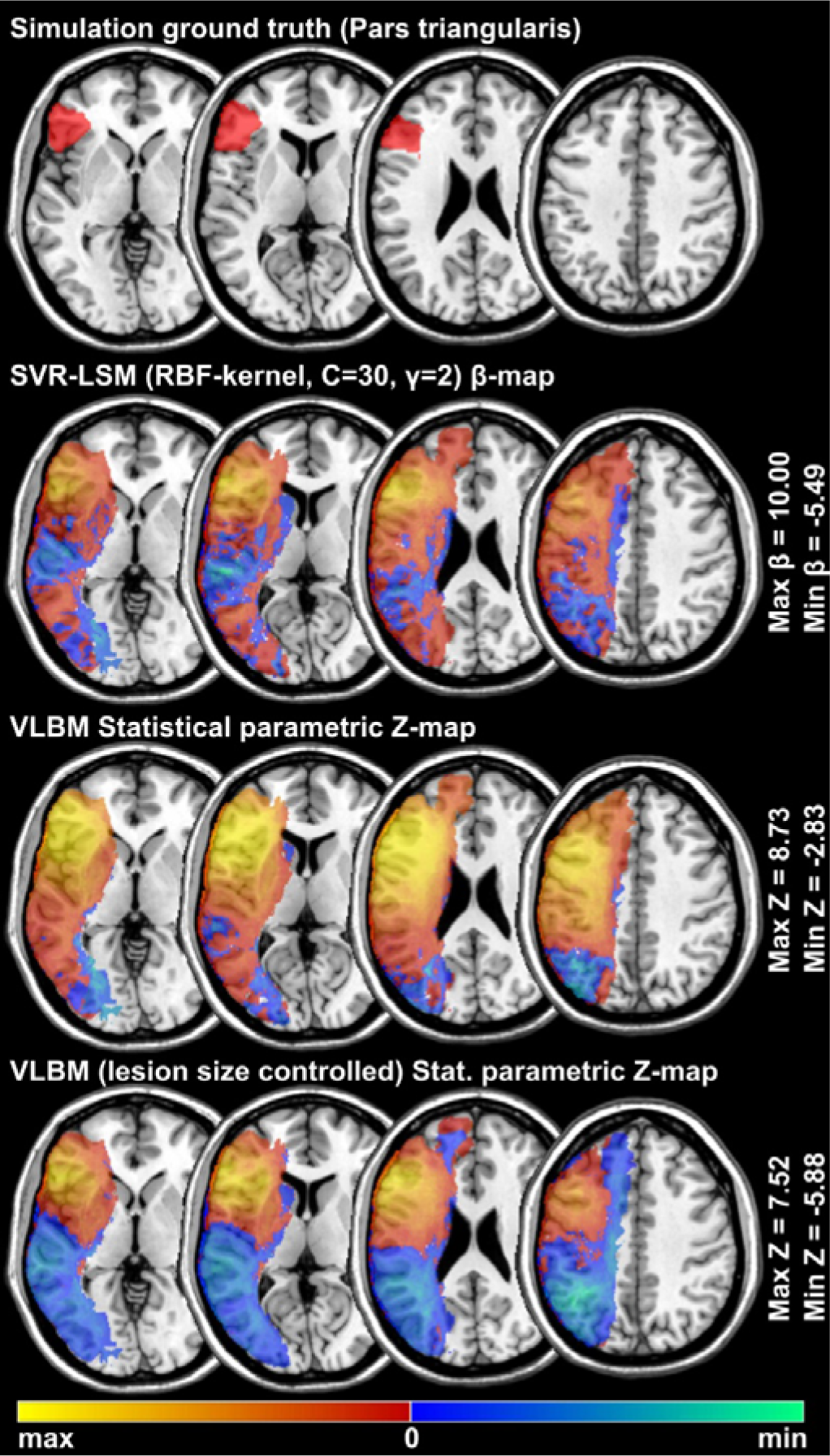
Positive and paradoxical negative signal in lesion-behaviour modelling. In the example lesion data set, a behavioural symptom was simulated for each patient based on damage to the pars triangularis. Thus, the only *causal* lesion-behaviour relation in the data is by definition a positive one. Regardless, both multivariate and mass-univariate lesion behaviour mapping not only find voxels with positive lesion-behaviour relation (red-yellow colour map), but also large areas with a considerably high paradoxical inverse lesion-behaviour relation (blue colour map), i.e. brain areas where damage is associated with a less severe symptom. Note that unthresholded maps are shown. Negative signal would surpass an uncorrected p-level of 0.05 in all analysis, however, in the present example it would not surpass a false discovery rate correction of q = 0.05.

### 4.2 Conclusion

Even in cases where no *causal* negative lesion-behaviour associations exist, negative lesion-behaviour associations might be present in the data. This effect and the lesion-anatomical biases of lesion behaviour mapping are two sides of the same coin: if damage to region A is causal for a deficit, an area B might also be positively associated with the deficit if regions A and B are often damaged together. This happens mainly in neighbouring regions along the vasculature towards the centres of the arterial territories (Mah et al., 2014; Sperber & Karnath, 2017; Sperber et al., 2019a). On the other hand, if the area A and another region area C are systematically *not* damaged together (e.g. because they are situated far from each other or in different arterial territories) damage in area C might be negatively associated with the deficit. While paradoxical negative signal in the present simulation example was lower than positive signal, uncorrected negative signal still surpassed common significance thresholds. Therefore, a proper correction for multiple comparisons is definitely necessary to prevent artefacts (see Karnath et al., 2018 for more information on multiple comparison correction). New lesion-deficit inference methods should be validated to only identify true (causal) negative lesion-deficit relations, and to be resistant to false alarms given non-causal negative lesion-deficit associations. Also, caution is advised if the behavioural target variable is controlled for other variables. It has been shown that such correction – if applied inappropriately – can enhance paradoxical lesion-deficit artefacts even above corrected thresholds (Sperber, Nolingberg, & Karnath, in press). Therefore, hypothesis-free interpretation of lesion-behaviour models does not provide an unambiguous view on paradoxical functional facilitation. This also holds true for multivariate models, that can yield large numbers of paradoxical lesion effects in real lesion behaviour data (see e.g. Toba et al., 2017). While lesion behaviour models are, in principle, powerful tools to identify possible paradoxical inversed lesion-deficit relations, we can find non-causal negative associations, and identification of functional facilitation in stroke should be conducted with caution in conjunction with a strong theoretical basis. A possible way to provide paradoxical lesion-deficit relations with a solid theoretical groundwork is to generate hypotheses based on single cases that show a possible post-stroke facilitation of symptoms (see Valero-Cabré et al., 2019), or on group studies in the case of enhancing lesion effects (Etcoff et al., 2000; Reverberi et al., 2005). A post-hoc validation option is the combination with other neuroscientific methods, such as the use of neuro-stimulation on regions identified in a VLBM analysis.

## 5 Different strategies for feature reduction in structural lesion data

In modern structural MR imaging, the brain can encompass over a million voxels. Models trained on voxel-wise information derived from such volumes thus can include over a million features. This massive number of features is a prime example for the curse of dimensionality, and it poses methodological challenges in any modelling process. A common procedure to overcome such an issue is feature reduction. Neighbouring voxels carry highly similar information. Therefore, information from multiple voxels could be merged into a single feature, and the overall number of features could be reduced. Accordingly, many studies that performed lesion-behaviour modelling – both for brain mapping and clinical outcome prediction – did so by merging voxel-wise data into region-wise data (Smith, Clithero, Rorden, & Karnath, 2013; Yourganov, Smith, Fridriksson, & Rorden, 2015; Zavaglia et al., 2015; Rondina et al., 2016; Yourganov, Fridriksson, Rorden, Gleichgerrcht, & Bonilha, 2016; Achilles et al., 2017; Toba et al., 2017; Hillis et al., 2018; Wilmskoetter et al., 2019). This is especially important in modelling algorithms that can handle only a limited number of features due to mathematical or computational restrictions (Smith et al., 2013; Zavaglia et al., 2015; Toba et al., 2017). Regions in these studies were defined by brain atlases which parcellate the brain based on, e.g., functional (Faria et al., 2012; Joliot et al., 2015; Glasser et al., 2016) or morphological (Tzourio-Mazoyer et al., 2002) criteria. Importantly, in order not to lose any data in the process of feature reduction, features have to be meaningfully merged. To do so in the modelling of brain imaging data, we aim to define brain areas in a way that minimises differences between voxels inside an area, while it maximises differences between voxels in different areas. In other words, a good parcellation aims to find homogeneous brain regions that are different from each other. If the parcellation of stroke imaging data is not carried out in a meaningful way, voxel-based analyses might easily outperform any region-based analysis (see e.g. Rondina et al., 2016).

An intuitive strategy to meaningfully merge voxels into regions is to use a brain parcellation based on functional data. Averaging data inside a functional module is a common approach when analysing functional imaging data (Poldrack, 2007), and the same can be done with lesion data (Pustina et al., 2018). The rationale is that damage to different parts of a homogeneous functional unit has equal effects on brain functionality. Damage to voxels inside this functional unit can thus be represented by only a single variable, such as the proportion of damaged voxels inside this region.

However, besides functional parcellation, a second strategy to meaningfully parcellate the brain exists in lesion data modelling. At this point, the lesion-anatomical dependence between brain regions comes back into play: the damage status of all brain regions is more or less correlated. Directly neighbouring voxels are often damaged or not damaged together. Therefore, the resolution of anatomical lesion data in group studies is smaller than the whole imaging volume, and some software tools for lesion behaviour mapping, by default, merge voxels that carry perfectly correlating anatomical information (Rorden et al., 2007; Pustina et al., 2018). One could now go even further and merge not only voxels that perfectly correlate, but also voxels that highly correlate. This could be done, e.g., by dimension reduction techniques such as principal component analysis (e.g. Moulton, Valabregue, Lehéricy, Samson, & Rosso, 2019; Loughnan et al., in press). The rationale for such lesion-anatomical dimension reduction is that we merge voxels whose relation to the behavioural variable cannot be further differentiated anyway, because the damage status of these voxels correlates too highly in a patient sample.

There are compelling reasons for both functional and lesion-anatomical parcellation. A conflict might now arise when functional brain areas and typical lesion anatomy differ. In such case, optimising a parcellation according to one strategy might hamper the parcellation quality when looking through the lens of the other strategy. For example, a functional parcellation of two brain areas might be worthless if we apply it on imaging of a stroke patient sample in which all patients either have damage to both regions or to neither of both.

### 5.1 Simple example

To illustrate the issue, Figure 4 shows two different pairs of voxels. These voxels have been manually chosen to best depict how functional and lesion-anatomical parcellation can markedly differ. Functional similarity was defined by reference to a multi-modal atlas based on different task-related and resting-state functional measures as well as myelin maps and cortical thickness (Glasser et al., 2016), which was converted into a volumetric parcellation (Pustina et al., 2018). Lesion-anatomical similarity between the binary lesion status of both voxels in the example lesion sample was defined by Pearson’s φ coefficient, which is a correlation measure for two dichotomous variables.

**Figure 4.**
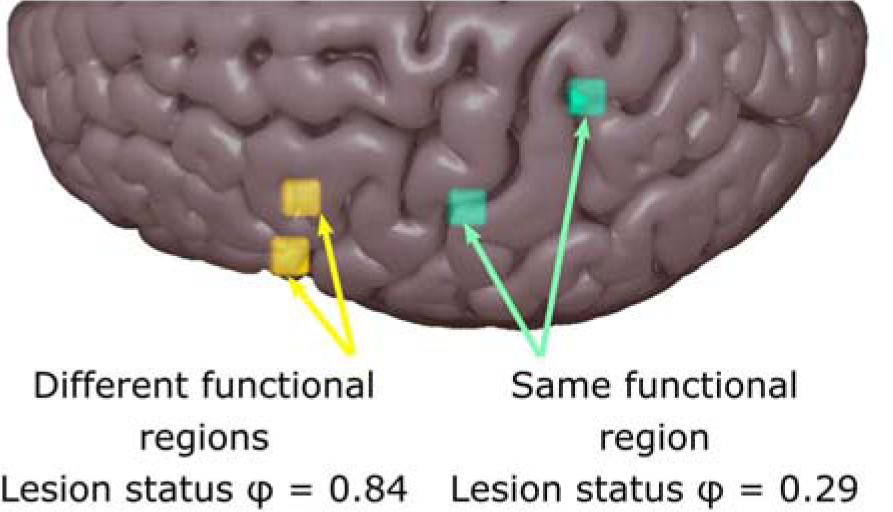
Functional and lesion-anatomical (dis-)similarity of voxels. Two pairs of voxels are chosen to illustrate possible conflicts between functional and lesion-anatomical parcellation. The yellow voxels are chosen to originate from different functional areas (xyz = −56,16,21 parcel index 74 (see Glasser et al., 2016 supplementary); xyz = - 44,13,15 parcel index 108 frontal opercular area 4); the turquois voxels to originate from the same functional area (xyz = −23, −42, 60; xyz = −46, −19, 41; both parcel index 52). The similarity of lesion status in these voxels in the sample of 131 left hemisphere lesions, as given by Pearson’s φ, is diametrical to their functional similarity (all coordinates are in MNI-space).

### 5.2 Conclusion

Two different strategies are available for meaningful feature reduction of structural lesion data in lesion-behaviour modelling – functional and lesion-anatomical parcellation. As seen in the example, both strategies can lead to different solutions for voxels. Thus, simply maximising feature reduction by a single parcellation strategy might fail to maximise the fit of lesion-behaviour models. Careful fine-tuning might be required to achieve this goal, and sample characteristics and size can affect lesion anatomy, possibly requiring to some degree individualized, data-driven solutions in every new sample. Further, it will be relevant what we want to achieve with the models. If we aim to investigate the individual role of anatomical features in such models – as done in lesion-behaviour mapping – it is essential that the features still resemble actual anatomical entities that allow us to obtain interpretable results. This, however, is not required if we only aim to maximise model fit between behaviour and anatomical data, as in stroke outcome prediction.

## 6 Model fit in lesion behaviour modelling and non-topographical features

A key aspect of causal relations in lesion-behaviour modelling has been omitted in the previous sections: it is not lesion location alone that causes a deficit, but an interplay of lesion location and many other factors that induce a deficit (for reviews see Price et al., 2017; Umarova, 2017). Factors such as age, neuropsychological co-morbidity, or pre-morbid cognitive status can affect the initial severity of a symptom, and even the more they will affect how well a patient can recover in the long term. Therefore, modelling of post-stroke deficits only based on topographical lesion information might, in many situations, be deemed to only achieve mediocre prediction performance. This effect is paralleled by sources of uncontrollable noise in the data, such as inter-individual anatomical differences, or noise in the processing of imaging data or in the assessment of the behavioural target variable (see Figure 5 for a schematic overview). This comes with important implications depending on the intention behind modelling the behavioural target variable based on anatomical lesion information.

**Figure 5.**
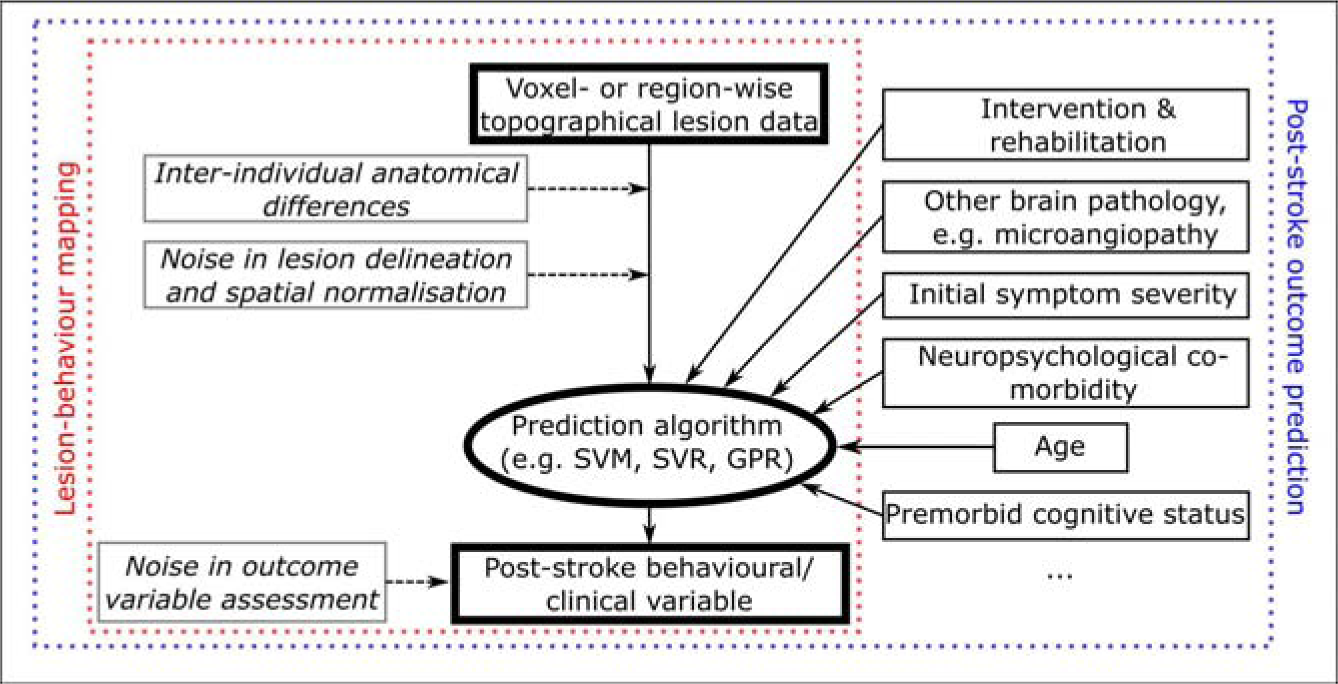
Possibly relevant variables in lesion-behaviour modelling. A schematic of non-topographical variables that might be of relevance in lesion behaviour modelling. Non-observable variables that introduce noise in lesion behaviour modelling, and thus generally hamper model fit, are shown in italic and with dashed lines. The red box shows the common variable set in lesion behaviour mapping, where non-topographical variables are commonly ignored. This can further limit model fit, even if the algorithm properly captures the variance explainable by the lesion topography.

While the modelling procedure can principally be the same both in post-stroke outcome prediction and in lesion-behaviour mapping, the criteria when evaluating model quality are different. Model quality in post stroke outcome prediction is principally straight forward to evaluate – a good model needs to be generalisable with a high, clinically relevant model fit to the population. This means that a model needs to provide good predictions in a previously unknown sample, i.e. the so-called test dataset. The criteria to access model quality are different in lesion-behaviour mapping, when the role of individual features is interpreted. Besides model fit, generalisability of feature weights respectively of topographical brain-lesion inference is desired (Rasmussen, Hansen, Madsen, Churchill, & Strother, 2012). Therefore, if the same behavioural variable is mapped in two independent data samples, the topographical inference should highly correspond between both analyses, and, in the case of e.g. SVR-LSM (Zhang et al., 2014), voxel-wise parameter weights in both should correlate. This gives rise to different implications for post-stroke outcome prediction and lesion-behaviour mapping. In post-stroke outcome prediction, a requirement might be to include some non-imaging parameters to obtain a model that can make good predictions. In lesion-behaviour mapping, we can include such variables into statistical models, and we can identify variables that should or should not be controlled for by looking at the causal relations between variables (Sperber et al., in press). If we decide not to include such variables into the model, we might end up with only a low to mediocre model fit. This does not necessarily invalidate the method. In the end, the aim is to obtain replicable, valid lesion-deficit inference. If a model handles the signal and the noise such that it effectively captures the variance explained by topographical lesion information, this might suffice – even if there is substantial amount of unexplained variance remaining. To conclude, lesion-behaviour mapping analyses should not (only) be judged by their model fit.

## 7 Conclusion and perspective

By reframing ‘causality’ in lesion-deficit inference, it can be shown that the validity of lesion behaviour mapping is affected by lesion anatomy, and that neither univariate nor multivariate algorithms can fully overcome this issue. While this is definitely a methodological limitation to keep in mind, I believe that the significance of this issue has been overstated. Not only did the present simulations not fully replicate an example that probably illustrated the worst-case shortcomings of voxel-wise lesion behaviour mapping, several previous studies have put the finding of a bias induced by lesion anatomy into a more nuanced perspective (Sperber & Karnath, 2017; Pustina et al., 2018). A strategy to counteract such bias could be to perform lesion-behaviour mapping in a more hypothesis-guided way, if possible. Results in voxel-wise lesion behaviour mapping could be critically interpreted from the perspective of a priori hypotheses. In the case of a strong hypothesis, one might even deviate from the whole-brain voxel-wise approach and instead only investigate regions of interest. It is also important to keep in mind that there is no neuroscientific gold standard method to map the human brain. All neuroscientific methods bear some limitations, and no single method provides us with perfect insights into functional brain anatomy. However, the available methods nicely complement each other (Rorden & Karnath, 2004).

Another issue of mass-univariate lesion behaviour mapping is the functional dependence between voxels, which could bias topographical results (Rorden et al., 2009; Mah et al., 2014). This issue can be resolved by using multivariate algorithms (Mah et al., 2014; Zhang et al., 2014; Pustina et al., 2018). Contrary to what has been stated elsewhere (Xu et al., 2018), this issue, and only this issue, is the main reason to use multivariate lesion behaviour mapping. Multivariate methods are still relatively new to the field, and many open questions on how to perform multivariate lesion-brain inference in the best manner are still open (see, e.g., DeMarco & Turkeltaub, 2018; Sperber et al., 2019a). Currently, two distinct main strategies to perform hypothesis-free voxel-wise lesion behaviour mapping were implemented and validated (Zhang et al., 2014; Pustina et al., 2018), and both have the potential to be further refined. Further, several region-based approaches (e.g. Yourganov et al., 2015; Achilles et al, 2017) also appear to be viable options for lesion-deficit inference. Thus, another open question is how to optimally include topographical lesion data into models, and under which conditions region-wise analysis can be superior to voxel-wise analysis.

In many cases, purely structural topographical lesion information is not sufficient to fully understand lesion-deficit relations. Focal damage can affect parts of structural and functional brain networks, and thus functional disruption might go beyond the actually lesioned area (Catani et al., 2012). Accordingly, lesion-behaviour modelling can be improved by looking beyond the mere topographies of structural damage. For example, functional or structural connectome imaging (e.g., Boes et al., 2015; Siegel et al., 2016; Kuceyeski et al., 2016; Gleichgerrcht, Fridriksson, Rorden, & Bonilha, 2017) or perfusion imaging (Karnath et al., 2005) can be used to better understand or predict lesion-deficit relations. Further, structural lesion information can be combined with, for example, functional or structural connectome data into models using multimodal imaging (Pustina et al., 2017; Foulon et al., 2018). Are such approaches a solution to overcome the causation vs. association issue in structural lesion-deficit relations? This is unlikely. Just as the damage status between two brain regions can non-causally be associated, abnormalities in functional networks or in fibre tract integrity might be non-causally associated, simply as the vasculature supplies brain tissue underlying these networks or fibre tracts in a systematic manner. To conclude, lesion-deficit inference based on any kind of connectome data is well-suited to complement VLBM and MLBM, but it does not overcome the main issues of lesion-behaviour modelling identified in the present article.

Stroke is the aetiology that is most commonly investigated by lesion behaviour mapping methods, yet focal brain damage induced by other aetiologies like brain tumours, surgical resections, or focal damage in inflammatory diseases is also sometimes investigated. Conclusions drawn in this article will likely hold true for all these aetiologies. Brain damage is always in some way systematic, and all aetiologies induce damage to one or multiple areas of connected brain voxels. However, the lesion-anatomical dependence between voxels will vary between aetiologies, depending on typical location, patterns and size of brain lesions.

The present paper’s most important implication for post-stroke outcome prediction is that feature selection and feature reduction are not intuitively straight-forward in lesion-behaviour models. Predictive features cannot only be found in brain regions where damage actually causes a long-lasting deficit, but in many areas in the brain. Another implication is that many features are highly redundant as their lesion-anatomical or functional information is highly dependent. Thus, it seems that to obtain excellent prediction algorithms, we have to embark on a long journey of refining modelling strategies while having in mind the high-dimensional complexity of lesion data.

## Abbreviations

VLBM: voxel-based lesion behaviour mapping
MLBM: multivariate lesion behaviour mapping
SVM: support-vector machine

## Acknowledgements

This work was supported by the Deutsche Forschungsgemeinschaft (KA 1258/23-1). I thank Hannah Tomczyk, Jonathan Glenday, and Daniel Wiesen for helpful comments on the manuscript. Declarations of interest: none.

## References

Achilles, E.I.S., Weiss, P.H., Fink, G.R., Binder, E., Price, C.J., Hope, T.M.H., 2017. Using multi-level Bayesian lesion-symptom mapping to probe the body-part-specificity of gesture imitation skills. Neuroimage 161, 94–103. https://doi.org/10.1016/j.neuroimage.2017.08.036

Adolphs, R., 2016. Human Lesion Studies in the 21st Century. Neuron 90, 1151–1153. https://doi.org/10.1016/j.neuron.2016.05.014

Bates, E., Wilson, S.M., Saygin, A.P., Dick, F., Sereno, M.I., Knight, R.T., Dronkers, N.F., 2003. Voxel-based lesion-symptom mapping. Nat. Neurosci. 6, 448–50. https://doi.org/10.1038/nn1050

Boes, A. D., Prasad, S., Liu, H., Liu, Q., Pascual-Leone, A., Caviness, V. S., & Fox, M. D. (2015). Network localization of neurological symptoms from focal brain lesions. Brain, 138, 3061–3075. https://doi.org/10.1093/brain/awv228

Carter, A.R., McAvoy, M.P., Siegel, J.S., Hong, X., Astafiev, S. V., Rengachary, J., Zinn, K., Metcalf, N. V., Shulman, G.L., Corbetta, M., 2017. Differential white matter involvement associated with distinct visuospatial deficits after right hemisphere stroke. Cortex 88, 81–97. https://doi.org/10.1016/j.cortex.2016.12.009

Catani, M., Dell’Acqua, F., Bizzi, A., Forkel, S. J., Williams, S. C., Simmons, A.,… Thiebaut de Schotten, M. (2012). Beyond cortical localization in clinico-anatomical correlation. Cortex, 48(10), 1262–1287. https://doi.org/10.1016/j.cortex.2012.07.001

Chang, C., Lin, C., 2011. LIBSVM. ACM Trans. Intell. Syst. Technol. 2, 1–27. https://doi.org/10.1145/1961189.1961199

DeMarco, A.T., Turkeltaub, P.E., 2018. A multivariate lesion symptom mapping toolbox and examination of lesion-volume biases and correction methods in lesion-symptom mapping. Hum. Brain Mapp. 39, 4169–4182. https://doi.org/10.1002/hbm.24289

Dickens, J.V., Fama, M.E., DeMarco, A.T., Lacey, E.H., Friedman, R.B., Turkeltaub, P.E., 2019. Localization of Phonological and Semantic Contributions to Reading. J. Neurosci. 39, 5361–5368. https://doi.org/10.1523/JNEUROSCI.2707-18.2019

Etcoff, N.L., Ekman, P., Magee, J.J., Frank, M.G., 2000. Lie detection and language comprehension. Nature 405, 139–139. https://doi.org/10.1038/35012129

Faria, A.V., Joel, S.E., Zhang, Y., Oishi, K., van Zjil, P.C.M., Miller, M.I., Pekar, J.J., Mori, S., 2012. Atlas-based analysis of resting-state functional connectivity: evaluation for reproducibility and multi-modal anatomy-function correlation studies. Neuroimage 61, 613–21. https://doi.org/10.1016/j.neuroimage.2012.03.078

Fecteau, S., Pascual-Leone, A., Théoret, H., 2006. Paradoxical Facilitation of Attention in Healthy Humans. Behav. Neurol. 17, 159–162. https://doi.org/10.1155/2006/632141

Foulon, C., Cerliani, L., Kinkingnéhun, S., Levy, R., Rosso, C., Urbanski, M., Volle, E., Thiebaut de Schotten, M., 2018. Advanced lesion symptom mapping analyses and implementation as BCBtoolkit. Gigascience 7, 1–17. https://doi.org/10.1093/gigascience/giy004

Gläscher, J., Adolphs, R., Tranel, D., 2019. Model-based lesion mapping of cognitive control using the Wisconsin Card Sorting Test. Nat. Commun. 10, 20. https://doi.org/10.1038/s41467-018-07912-5

Glasser, M.F., Coalson, T.S., Robinson, E.C., Hacker, C.D., Harwell, J., Yacoub, E., Ugurbil, K., Andersson, J., Beckmann, C.F., Jenkinson, M., Smith, S.M., Van Essen, D.C., 2016. A multi-modal parcellation of human cerebral cortex. Nature 536, 171–8. https://doi.org/10.1038/nature18933

Gleichgerrcht, E., Fridriksson, J., Rorden, C., Bonilha, L., 2017. Connectome-based lesion-symptom mapping (CLSM): A novel approach to map neurological function. NeuroImage Clin. 16, 461–467. https://doi.org/10.1016/j.nicl.2017.08.018

Hillis, A.E., Beh, Y.Y., Sebastian, R., Breining, B., Tippett, D.C., Wright, A., Saxena, S., Rorden, C., Bonilha, L., Basilakos, A., Yourganov, G., Fridriksson, J., 2018. Predicting recovery in acute poststroke aphasia. Ann. Neurol. 83, 612–622. https://doi.org/10.1002/ana.25184

Hope, T.M.H., Leff, A.P., Price, C.J., 2018. Predicting language outcomes after stroke: Is structural disconnection a useful predictor? NeuroImage Clin. 19, 22–29. https://doi.org/10.1016/j.nicl.2018.03.037

Howard, C.M., Smith, L.L., Coslett, H.B., Buxbaum, L.J., 2019. The role of conflict, feedback, and action comprehension in monitoring of action errors: Evidence for internal and external routes. Cortex 115, 184–200. https://doi.org/10.1016/j.cortex.2019.01.032

Inoue, K., Madhyastha, T., Rudrauf, D., Mehta, S., Grabowski, T., 2014. What affects detectability of lesion–deficit relationships in lesion studies? NeuroImage Clin. 6, 388–397. https://doi.org/10.1016/j.nicl.2014.10.002

Joliot, M., Jobard, G., Naveau, M., Delcroix, N., Petit, L., Zago, L., Crivello, F., Mellet, E., Mazoyer, B., Tzourio-Mazoyer, N., 2015. AICHA: An atlas of intrinsic connectivity of homotopic areas. J. Neurosci. Methods 254, 46–59. https://doi.org/10.1016/j.jneumeth.2015.07.013

Kapur, N., 1996. Paradoxical functional facilitation in brain-behaviour research. A critical review. Brain 119, 1775–90. https://doi.org/10.1093/brain/119.5.1775

Karnath, H.O., Zopf, R., Johannsen, L., Berger, M.F., Nägele, T., Klose, U., 2005. Normalized perfusion MRI to identify common areas of dysfunction: Patients with basal ganglia neglect. Brain 128, 2462–2469. https://doi.org/10.1093/brain/awh629

Karnath, H.-O., Sperber, C., Rorden, C., 2018. Mapping human brain lesions and their functional consequences. Neuroimage 165, 180–189. https://doi.org/10.1016/j.neuroimage.2017.10.028

Kuceyeski, A., Navi, B.B., Kamel, H., Raj, A., Relkin, N., Toglia, J., Iadecola, C., O’Dell, M., 2016. Structural connectome disruption at baseline predicts 6-months post-stroke outcome. Hum. Brain Mapp. 2601, 2587–2601. https://doi.org/10.1002/hbm.23198

Loughnan, R., Lorca-Puls, D.L., Gajardo-Vidal, A., Espejo-Videla, V., Gillebert, C.R., Mantini, D., Price, C.J., Hope, T.M.H, in press. Generalizing post-stroke prognoses from research data to clinical data. NeuroImage Clin.

Mah, Y.-H., Husain, M., Rees, G., Nachev, P., 2014. Human brain lesion-deficit inference remapped. Brain 137, 2522–31. https://doi.org/10.1093/brain/awu164

Moulton, E., Valabregue, R., Lehéricy, S., Samson, Y., Rosso, C., 2019. Multivariate prediction of functional outcome using lesion topography characterized by acute diffusion tensor imaging. NeuroImage Clin. 23, 101821. https://doi.org/10.1016/j.nicl.2019.101821

Nachev, P., 2015. The first step in modern lesion-deficit analysis. Brain 138, e354. https://doi.org/10.1093/brain/awu275

Pearl, J., Glymour, M., Jewell, N.P, 2016. Causal inference in statistics, first ed. Wiley, Chichester.

Pearl, J., Mackenzie, D., 2018. The book of why, first ed. Basic Books, New York.

Poldrack, R.A., 2007. Region of interest analysis for fMRI. Soc. Cogn. Affect. Neurosci. 2, 67–70. https://doi.org/10.1093/scan/nsm006

Price, C.J., Hope, T.M., Seghier, M.L., 2017. Ten problems and solutions when predicting individual outcome from lesion site after stroke. Neuroimage 145, 200–208. https://doi.org/10.1016/j.neuroimage.2016.08.006

Pustina, D., Avants, B., Faseyitan, O.K., Medaglia, J.D., Coslett, H.B., 2018. Improved accuracy of lesion to symptom mapping with multivariate sparse canonical correlations. Neuropsychologia 115, 154–166. https://doi.org/10.1016/j.neuropsychologia.2017.08.027

Pustina, D., Coslett, H.B., Ungar, L., Faseyitan, O.K., Medaglia, J.D., Avants, B., Schwartz, M.F., 2017. Enhanced estimations of post-stroke aphasia severity using stacked multimodal predictions. Hum. Brain Mapp. 38, 5603–5615. https://doi.org/10.1002/hbm.23752

Rasmussen, P.M., Hansen, L.K., Madsen, K.H., Churchill, N.W., Strother, S.C., 2012. Model sparsity and brain pattern interpretation of classification models in neuroimaging. Pattern Recognit. 45, 2085–2100. https://doi.org/10.1016/j.patcog.2011.09.011

Reverberi, C., Toraldo, A., D’Agostini, S., Skrap, M., 2005. Better without (lateral) frontal cortex? Insight problems solved by frontal patients. Brain 128, 2882–2890. https://doi.org/10.1093/brain/awh577

Rondina, J.M., Filippone, M., Girolami, M., Ward, N.S., 2016. Decoding post-stroke motor function from structural brain imaging. NeuroImage. Clin. 12, 372–80. https://doi.org/10.1016/j.nicl.2016.07.014

Rondina, J.M., Park, C.H., Ward, N.S., 2017. Brain regions important for recovery after severe post-stroke upper limb paresis. J. Neurol. Neurosurg. Psychiatry 88, 737–743. https://doi.org/10.1136/jnnp-2016-315030

Rorden, C., Karnath, H.-O., 2004. Using human brain lesions to infer function: a relic from a past era in the fMRI age? Nat. Rev. Neurosci. 5, 813–9. https://doi.org/10.1038/nrn1521

Rorden, C., Karnath, H.-O., Bonilha, L., 2007. Improving lesion-symptom mapping. J. Cogn. Neurosci. 19, 1081–8. https://doi.org/10.1162/jocn.2007.19.7.1081

Rorden, C., Fridriksson, J., Karnath, H.-O., 2009. An evaluation of traditional and novel tools for lesion behavior mapping. Neuroimage 44, 1355–62. https://doi.org/10.1016/j.neuroimage.2008.09.031

Siegel, J.S., Ramsey, L.E., Snyder, A.Z., Metcalf, N. V, Chacko, R. V, Weinberger, K., Baldassarre, A., Hacker, C.D., Shulman, G.L., Corbetta, M., 2016. Disruptions of network connectivity predict impairment in multiple behavioral domains after stroke. Proc. Natl. Acad. Sci. U. S. A. 113, E4367–76. https://doi.org/10.1073/pnas.1521083113

Smith, D. V, Clithero, J.A., Rorden, C., Karnath, H.-O., 2013. Decoding the anatomical network of spatial attention. Proc. Natl. Acad. Sci. U. S. A. 110, 1518–23. https://doi.org/10.1073/pnas.1210126110

Sperber, C., Karnath, H.-O., 2017. Impact of correction factors in human brain lesion-behavior inference. Hum. Brain Mapp. 38, 1692–1701. https://doi.org/10.1002/hbm.23490

Sperber, C., Karnath, H.-O., 2018. On the validity of lesion-behaviour mapping methods. Neuropsychologia 115, 17–24. https://doi.org/10.1016/j.neuropsychologia.2017.07.035

Sperber, C., Wiesen, D., Karnath, H., 2019a. An empirical evaluation of multivariate lesion behaviour mapping using support vector regression. Hum. Brain Mapp. 40, 1381–1390. https://doi.org/10.1002/hbm.24476

Sperber, C., Wiesen, D., Goldenberg, G., & Karnath, H.-O. 2019b. A network underlying human higher-order motor control: Insights from machine learning-based lesion-behaviour mapping in apraxia of pantomime. Cortex. 121, 308–321. https://doi.org/10.1016/j.cortex.2019.08.023

Sperber, C., Nolingberg, C., Karnath, H., in press. Post-stroke cognitive deficits rarely come alone: handling co-morbidity in lesion behaviour mapping. Hum. Brain Mapp. https://doi.org/10.1002/hbm.24885

Sprague, J.M., 1966. Interaction of cortex and superior colliculus in mediation of visually guided behavior in the cat. Science. 153, 1544–1547. https://doi.org/10.1126/science.153.3743.1544

Thye, M., Mirman, D., 2018. Relative contributions of lesion location and lesion size to predictions of varied language deficits in post-stroke aphasia. NeuroImage Clin. 20, 1129–1138. https://doi.org/10.1016/j.nicl.2018.10.017

Toba, M.N., Zavaglia, M., Rastelli, F., Valabrégue, R., Pradat-Diehl, P., Valero-Cabré, A., Hilgetag, C.C., 2017. Game theoretical mapping of causal interactions underlying visuospatial attention in the human brain based on stroke lesions. Hum. Brain Mapp. 38, 3454–3471. https://doi.org/10.1002/hbm.23601

Tzourio-Mazoyer, N., Landeau, B., Papathanassiou, D., Crivello, F., Etard, O., Delcroix, N., Mazoyer, B., Joliot, M., 2002. Automated anatomical labeling of activations in SPM using a macroscopic anatomical parcellation of the MNI MRI single-subject brain. Neuroimage 15, 273–289. https://doi.org/10.1006/nimg.2001.0978

Umarova, R.M., 2017. Adapting the concepts of brain and cognitive reserve to post-stroke cognitive deficits: Implications for understanding neglect. Cortex. 97, 327–338. https://doi.org/10.1016/j.cortex.2016.12.006

Valero-Cabré, A., Toba, M.N., Hilgetag, C.C., Rushmore, R.J., 2019. Perturbation-driven paradoxical facilitation of visuo-spatial function: Revisiting the “Sprague effect”. Cortex. https://doi.org/10.1016/j.cortex.2019.01.031

Wilmskoetter, J., Fridriksson, J., Gleichgerrcht, E., Stark, B.C., Delgaizo, J., Hickok, G., Vaden, K.I., Hillis, A.E., Rorden, C., Bonilha, L., 2019. Neuroanatomical structures supporting lexical diversity, sophistication, and phonological word features during discourse. NeuroImage Clin. 24, 101961. https://doi.org/10.1016/j.nicl.2019.101961

Wong, A.L., Jax, S.A., Smith, L.L., Buxbaum, L.J., Krakauer, J.W., 2019. Movement Imitation via an Abstract Trajectory Representation in Dorsal Premotor Cortex. J. Neurosci. 39, 3320–3331. https://doi.org/10.1523/JNEUROSCI.2597-18.2019

Xu, T., Jha, A., Nachev, P., 2018. The dimensionalities of lesion-deficit mapping. Neuropsychologia 115, 134–141. https://doi.org/10.1016/j.neuropsychologia.2017.09.007

Yourganov, G., Smith, K.G., Fridriksson, J., Rorden, C., 2015. Predicting aphasia type from brain damage measured with structural MRI. Cortex. 73, 203–15. https://doi.org/10.1016/j.cortex.2015.09.005

Yourganov, G., Fridriksson, J., Rorden, C., Gleichgerrcht, E., Bonilha, L., 2016. Multivariate Connectome-Based Symptom Mapping in Post-Stroke Patients: Networks Supporting Language and Speech. J. Neurosci. 36, 6668–6679. https://doi.org/10.1523/JNEUROSCI.4396-15.2016

Zavaglia, M., Forkert, N.D., Cheng, B., Gerloff, C., Thomalla, G., Hilgetag, C.C., 2015. Mapping causal functional contributions derived from the clinical assessment of brain damage after stroke. NeuroImage Clin. 9, 83–94. https://doi.org/10.1016/j.nicl.2015.07.009

Zhang, Y., Kimberg, D.Y., Coslett, H.B., Schwartz, M.F., Wang, Z., 2014. Multivariate lesion-symptom mapping using support vector regression. Hum. Brain Mapp. 5876, 5861–5876. https://doi.org/10.1002/hbm.22590

Zhao, L., Biesbroek, J.M., Shi, L., Liu, W., Kuijf, H.J., Chu, W.W.C., Abrigo, J.M., Lee, R.K.L., Leung, T.W.H., Lau, A.Y.L., Biessels, G.J., Mok, V., Wong, A., 2018. Strategic infarct location for post-stroke cognitive impairment: A multivariate lesion-symptom mapping study. J. Cereb. Blood Flow Metab. 38, 1299–1311. https://doi.org/10.1177/0271678X17728162

